# Fibrosis burden and spatial pattern affect left atrial appendage stasis in multi-beat multi-physics simulations

**DOI:** 10.64898/2026.09.26.754330

**Authors:** Yvonne Stöcker, Christoph M. Augustin, Manuel Guerrero-Hurtado, Oscar Flores, Alejandro Gonzalo, Åshild Telle, Patrick M. Boyle, Nazem Akoum, Juan C. del Alamo, Manuel García-Villalba

## Abstract

**Purpose:** Atrial fibrosis is associated with thromboembolic risk in atrial fibrillation, but it remains unclear how fibrosis-related contractile impairment alters blood transport in the left atrium (LA) and left atrial appendage (LAA). Prior multi-physics simulations coupling electromechanical (EM) and computational fluid dynamics (CFD) models showed that fibrosis impairs LA function and flow, but were limited to booster-only contraction and could not separate the effects of fibrosis burden from its spatial distribution. Here, we examined how fibrosis burden and spatial pattern independently relate to long-term LA and LAA hemodynamics in full-cycle, multi-beat simulations on a common LA anatomy.

**Methods:** Three fibrosis patterns derived from late gadolinium enhancement MRI were applied globally to a common LA anatomy at matched burdens of 15.6% and 31.2%, yielding six global-fibrosis cases and a non-fibrotic reference. Two patterns were also confined to the LAA at local burdens of 10%, 25%, and 50%, yielding six LAA-fibrosis cases. A 3D EM model of the LA anatomy coupled to a 0D circulatory model resolved the reservoir, conduit, and booster phases, and the EM-derived wall motion drove subsequent multi-beat CFD simulations. Region-specific emptying fraction (EF), non-dimensional kinetic energy (KE), and the 90th percentile of blood residence time (T*_R_*) were evaluated in the LA and LAA.

**Results:** Left ventricular stroke volume remained nearly constant across cases, whereas increasing global fibrosis burden progressively reduced LA and LAA EF (LA: 36% to 30–31%; LAA: 44% to 28–38%), lowered both active and passive LA EF, and monotonically increased LA T*_R_* across all patterns. Non-dimensional LA KE showed an approximately quadratic association with LA EF across patterns, whereas LAA KE was more scattered but still increased with LAA EF. In contrast, the 90th percentile of LAA T*_R_* did not increase monotonically with decreasing LAA EF for one pattern. When fibrosis was confined to the LAA, non-dimensional LAA KE collapsed onto a single relationship with LAA EF for both patterns, whereas LAA T*_R_* followed distinct, non-monotonic trends.

**Conclusions:** Fibrosis burden, through its association with functional impairment, predominantly governed whole-LA hemodynamics, whereas LAA stasis additionally depended on the spatial distribution of fibrosis. In the LAA-fibrosis cases, LAA EF was a robust descriptor of chamber-averaged KE regardless of fibrosis pattern, but did not fully explain LAA residence time. These findings support evaluating imaging-derived fibrosis pattern, in addition to fibrosis burden and functional metrics, as a potential determinant of LAA transport and stasis. Its incremental value for thrombogenic-risk assessment requires prospective clinical validation.

## 1 Introduction

Ischemic stroke risk is elevated in patients with atrial fibrillation (AF), and current clinical tools used to guide anticoagulation, such as the CHA_2_DS_2_-VASc score, have only modest predictive performance. This limitation has motivated interest in additional patient-specific markers linked to the left atrium (LA) itself [1, 2]. In particular, structural and functional atrial characteristics may provide complementary information about thromboembolic risk beyond clinical comorbidity scores [2–4].

Atrial fibrosis, a hallmark of atrial myocardial remodeling, can be assessed non-invasively by late gadolinium enhancement (LGE)–magnetic resonance imaging (MRI) [5]. Greater atrial fibrosis, reflected by more extensive LGE, has been associated with impaired atrial reservoir, conduit, and booster function assessed by strain imaging [6–8]. In patients with AF, higher fibrosis burden has been associated with a history of stroke or transient ischemic attack [3, 9, 10] and with spontaneous echo contrast and left atrial appendage (LAA) thrombus on transesophageal echocardiography [4]. Patients with embolic stroke of undetermined source (ESUS), without a diagnosis of AF, have been found to have greater atrial fibrosis burden [11–13] and reduced LA strain [14]. Although these observational findings link atrial fibrosis to thromboembolic phenomena, they do not establish causality or identify the mechanisms underlying this association.

Fibrosis may contribute to thrombogenic conditions through at least two pathways. First, fibrosis-related impairment of atrial contraction may reduce blood motion and thereby promote stasis [4, 15, 16]. Second, fibrosis-related tissue injury may affect the coagulation cascade through mechanisms not captured by blood transport alone [4]. Consistent with a mechanical contribution, reduced LAA flow velocity and enlarged LAA orifice area have each been associated with increased stroke risk [17, 18].

Computational modeling can help isolate relationships among fibrosis, atrial function, and hemodynamics, but only a small number of computational fluid dynamics (CFD) studies have examined fibrosis in relation to intracardiac flow. Paliwal *et al.* [19] mapped fibrosis from LGE–MRI onto CFD simulations with prescribed wall motion and reported lower wall shear stress (WSS) in fibrotic than in non-fibrotic wall regions. The simulations did not explicitly model fibrosis-induced changes in wall motion. Thus, the reported WSS values probably reflect the local anatomy of fibrotic regions rather than fibrotic alterations in atrial mechanics. Combining rigid-wall CFD simulations with electro-anatomical maps and LGE–MRI, Adamopoulos *et al.* [20] reported regional co-localization of fibrosis or electrical scarring with higher time-averaged wall shear stress (TAWSS). At the whole-chamber scale and using image-derived wall motion, Parker *et al.* [21] found that global TAWSS decreased with increasing global fibrosis burden. These apparently diverging trends are not necessarily inconsistent, because the studies examined different hemodynamic quantities and spatial scales using distinct study designs and modeling assumptions. Nevertheless, the limited and heterogeneous literature leaves substantial uncertainty regarding how fibrosis affects atrial flow and blood stasis.

Gonzalo *et al.* [22] used a multi-physics computational setup that coupled fibrosis-related electromechanics (EM) changes with CFD simulations of LA blood flow. In their framework, an EM solver simulated contraction of four patient-specific LA anatomies, each with its own fibrosis burden and spatial distribution. The study reported that the hemodynamic effects of fibrosis were not limited to near-wall flow and that global flow kinetic energy correlated with global chamber function. For the LAA, however, stasis could not be explained by functional impairment alone, possibly reflecting additional roles of LAA morphology, interaction with flow from the LA body, or the spatial distribution of fibrotic tissue. However, because anatomy, fibrosis burden, and spatial distribution varied simultaneously among the cases, their respective contributions could not be isolated. Moreover, the simulation setup focused on the LA booster function, neglecting the reservoir and conduit phases. Thus, full-cycle and long-term blood transport could not be assessed.

The present study addresses these gaps by modeling fibrosis-related EM changes while systematically varying fibrosis burden and spatial distribution on a common LA anatomy. Full-cycle EM simulations incorporating a 0D model of the circulatory system provide wall motion and boundary conditions for subsequent multi-beat CFD simulations. Using this framework, we aim to examine how fibrosis burden and spatial distribution affect long-term LA and LAA blood transport. Blood residence time T*_R_*, a transport-based metric that quantifies the persistence of blood within a region, is used here to characterize blood stasis [23].

## 2 Methodology

### 2.1 Patients and Imaging

Atrial geometries and fibrosis distributions from three AF patients, denoted A, B, and C, were obtained from the cohort presented in Telle *et al.* [24]. To isolate the effects of fibrosis burden and spatial distribution from those of anatomy, all simulations used the geometry of patient A, while all three patients provided the fibrosis patterns. The patients were recruited at the University of Washington Medical Center and were scheduled for ablation. The acquisition and use of patient imaging data were approved by the Institutional Review Board of the University of Washington (STUDY00015081) and all participants provided informed consent for the collection and use of their data. LGE–MRI scans were acquired pre-ablation at the end of atrial diastole, providing the anatomy and spatial distributions of fibrotic tissue depicted in Figure 1. The corresponding total fibrotic burdens in the whole LA and in the LAA are reported in Table 1.

**Fig. 1.**
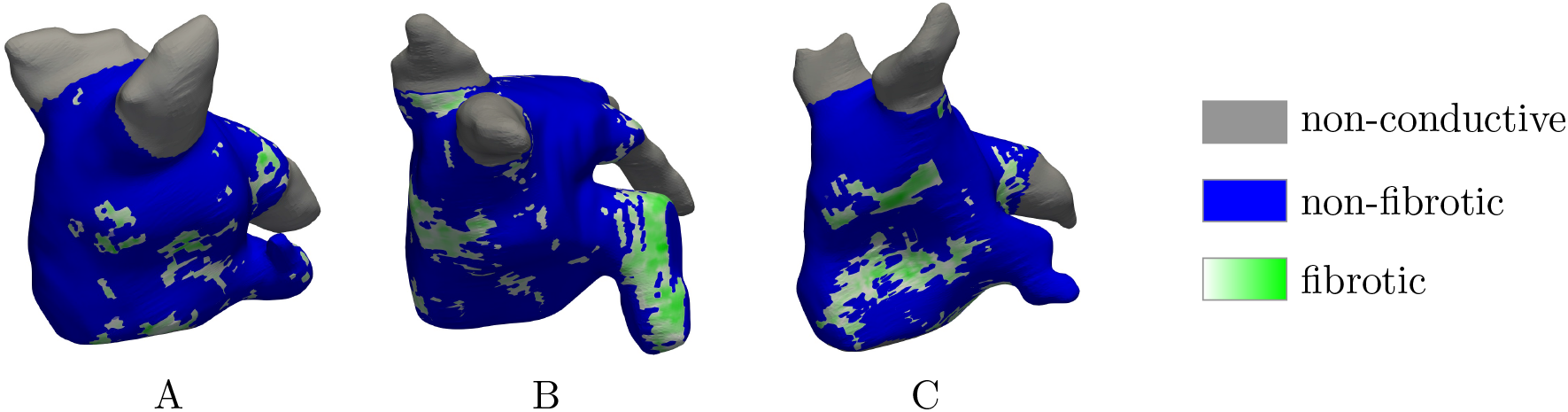
LGE maps showing the normalized LGE intensity of the fibrotic tissue for subjects A, B, and C

**Tab. 1.** Fibrotic burden [%] in the whole LA and LAA measured from the LGE–MRI scans.

| subject | A | B | C |
| --- | --- | --- | --- |
| LA | 15.6 | 23.9 | 17.9 |
| LAA | 6.88 | 39.57 | 12.43 |

### 2.2 Meshing for EM Simulations

Anatomy A was used for all the simulations in this study. The starting point for the EM mesh was the corresponding 3D triangulated endocardial surface mesh. To create a volumetric tetrahedral mesh of the myocardial tissue, the surface was extruded by 0.8 mm to generate the endocardial layer and by a further 1.2 mm to generate the epicardial layer. The average tetrahedral edge length was 500 µm. Auxiliary volumetric mesh elements were added to close the mitral valve (MV) and pulmonary vein (PV) orifices. These caps help preserve the physiological shape of the orifices and facilitate boundary-condition enforcement in both the EM and CFD simulations. Myocyte fiber orientation was assigned with a rule-based method described previously [25, 26].

### 2.3 Fibrosis Mapping and Case Design

The LGE maps shown in Figure 1 were used as three spatial patterns, each containing normalized LGE intensity values ranging from 0 to 1, with higher values indicating greater enhancement. First, the LGE distributions of subjects B and C were mapped onto the endocardial surface mesh of anatomy A using universal atrial coordinates [27] and subsequently extended through the wall thickness. Fibrosis was then assigned to the volumetric mesh as described by Telle *et al.* [24]: The volumetric elements were ranked by their mapped LGE intensity and assigned as fibrotic until a target burden was reached. To isolate the effect of spatial fibrosis distribution from that of total burden, all three patterns were applied at two matched global fibrosis burdens. The low-burden target was 15.6%, corresponding to the patient-specific burden of subject A (Tab. 1), whereas the high-burden target was 31.2%, obtained by doubling this value. Both target burdens were calculated from the volumetric elements, excluding the PVs and the artificial MV and PV caps.

Additionally, a non-fibrotic reference case, denoted nofib, was simulated on the same anatomy A. In this case, all myocardial elements were assigned non-fibrotic properties, giving zero fibrosis burden in both the LA and LAA.

Matching global LA fibrosis burden across patterns meant that local LAA burden could not be matched simultaneously. Motivated by this limitation, a second case set was generated, in which fibrotic tissue was confined to the LAA. This allowed for systematic variation in local appendage impairment while minimizing changes in global LA function. Patterns A and B were used for these LAA-fibrosis cases. Each pattern was applied with local LAA fibrosis burdens of 10%, 25%, and 50%. While the LGE data for pattern B permitted generating burdens up to 80% within the LAA, pattern A permitted a maximum burden of 24% using the assignment method described above. To overcome this limit, pattern-A target burdens exceeding 24% were generated by progressively dilating fibrotic regions: adjacent elements were assigned as fibrotic until the target burden was reached.

To characterize the spatial structure of the fibrosis patterns, fibrotic regions were decomposed into patches in the tetrahedral volume mesh. A patch was defined as a connected set of fibrotic tetrahedra linked through shared faces. For each global-fibrosis case, we report the number of patches N_patch_ and the arithmetic mean and maximum patch volume.

### 2.4 EM Simulations

The EM simulations were performed with the in-house code *CARPentry* [28, 29]. Details on the EM model, including baseline model parameters, are described elsewhere [22, 24, 30]. Here, we summarize the main features and the modifications applied to fibrotic tissue.

On the cell scale, electrophysiology (EP) was represented by the human atrial action-potential model of Courtemanche *et al.* [31] with modifications described by Bayer *et al.* [32], while the electrical activation was propagated through the myocardium on the macroscopic scale with a reaction-eikonal model [33]. Active stress was generated using a cell-level contraction model for human atria developed by Land and Niederer [34], with the maximum active tension T*_a_* scaled to 50 kPa in the non-fibrotic tissue. Passive stiffness was represented by a reduced Holzapfel–Ogden formulation with fiber dispersion using the parameters a = 2.92 kPa, b = 5.6, a*_f_* = 11.84 kPa, b*_f_* = 17.95, δ*_f_* = 0.09 for normal tissue [22, 30]. The artificial PV and MV caps were represented by a stiff Demiray material [24, 35].

In fibrotic elements, mechanical remodeling was represented by increasing the parameters a and a*_f_* in the model of passive stiffness fivefold [36] and scaling active tension by 0.5 [37]. Conduction velocities in fibrotic regions were reduced by 12%, consistent with the reduction reported in a recent study [38].

Atrial motion was constrained with Robin-type boundary conditions. Specifically, omnidirectional springs were applied to the PV caps, and normal springs were imposed on the epicardial surface. The interaction between the 3D EM model and the remaining parts of the circulatory system was incorporated by coupling it to the 0D lumped-parameter model *CircAdapt* [39, 40]. This model provided volumetric flow rates through the PVs and the MV (Q^PV^ and Q^MV^, respectively), and hydrostatic pressures, which were used to derive physiological pressure boundary conditions and volume constraints for the 3D LA structure [29]. *CircAdapt* was also used to impose a traction boundary condition on the MV annulus depending on ventricular contraction, thereby modeling atrioventricular plane displacement [24].

This setup differs from that of Gonzalo *et al.* [22], in which the LA contracted against a constant pressure of 10 mmHg, so that only the booster phase was represented. Here, the coupling to *CircAdapt* allowed the LA to additionally fill from the PVs during ventricular systole (reservoir phase) and to empty passively after MV opening (conduit phase). The simulations therefore resolved the full atrial cycle. The unloaded reference configuration of each case was determined as described previously [30, 41].

At the beginning of each heartbeat, electrical activation was triggered in the 3D EM model at three points near the right superior PV. These locations are the approximate activation sites observed for patient A [24].

All cases were simulated for 30 EM cycles at a heart rate of 60 bpm to ensure periodicity. To reduce computational cost, the mechanical components used a coarse time step of Δt_mech_ = 1 ms during the first 25 cycles. The final five cycles were performed with a refined time step of Δt_mech_ = 50 µs, matching the CFD time step described below. Throughout all cycles, the time step size for the EP components was set to Δt_EP_ = 25 µs. Endocardial motion during the final heartbeat (30^th^ cycle) was used as input to the CFD pipeline.

### 2.5 CFD Simulations

The CFD simulations followed the methodology of Stöcker *et al.* [42]: The incompressible Navier–Stokes equations were solved with the in-house flow solver TUCANGPU [43] using a fractional-step method. Blood was modeled as a Newtonian fluid with kinematic viscosity ν = 0.04 cm^2^/s. Second-order central differences were used for spatial discretization of the triply periodic cubic computational domain with an edge length of 13 cm and 256 uniformly distributed grid points in each direction (Δx_CFD_ ≈ 0.51 mm, 16.8 million grid cells). A three-substep low-storage semi-implicit Runge–Kutta method was used for time integration. With a constant time step size Δt_CFD_ = 50 µs, the Courant–Friedrichs–Lewy number remained below 0.3 in all simulations.

The moving LA wall was represented with an immersed boundary method (IBM) [44]. In this technique, the LA wall is represented by a cloud of marker points placed in the fixed Eulerian fluid grid. These marker points corresponded to the centroids of the triangular elements of the endocardial LA surface in the EM mesh. Because the final EM cycles and the CFD simulations used the same time step, the wall deformation was transferred to the flow solver without temporal interpolation. The no-slip condition at the LA wall was enforced by including volumetric forces as a source term in the fluid momentum equations in the vicinity of the LA marker points. These IBM forces were evaluated by interpolating flow quantities between the fixed Eulerian grid of the fluid and the instantaneous marker-point locations.

Flow through the PVs was imposed by extruding the PV cap surfaces upstream to create buffer regions. In these regions, a uniform surface-normal velocity was imposed instead of a no-slip condition. The total PV inflow rate obtained from the 0D model during the EM simulations was split among the four PVs to yield equal velocity in each vein.

For marker points on the MV cap, the no-slip condition was enforced only when no outflow was prescribed (Q^MV^ = 0 mL/s). When Q^MV^ > 0, no IBM forcing was applied on these points.

A second layer of marker points was generated by duplicating each point on the LA wall and MV cap and shifting each duplicate by one grid-step size Δx_CFD_ outward along the normal direction of its corresponding triangular element. This second layer reduced residual mass flux (leakage) across the immersed boundary, an issue reported by several research groups [42, 45, 46].

Blood residence time, T*_R_*, was modeled as a passive scalar transported by the fluid velocity 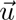 [47]:

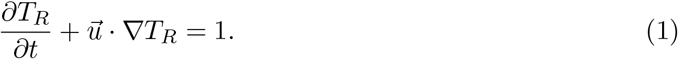

This equation was discretized in space with a third-order weighted essentially non-oscillatory (WENO) scheme [48], which provides numerical stability and accuracy in the absence of a diffusion term. In the buffer region upstream of the PV orifice caps described above, T*_R_* was set to zero using the IBM forcing procedure.

All CFD simulations were initialized from zero velocity and zero residence time. Running for 20 cardiac cycles ensured that T*_R_* growth saturated and reached a quasi-periodic state in all simulations. Then, 12 more cycles were simulated for post-processing.

### 2.6 Hemodynamic Metrics and Post-Processing

Hemodynamic quantities were evaluated separately in the LA and LAA. The LA region of interest (ROI) comprised the complete atrial blood volume, whereas the LAA ROI comprised only the blood volume inside the appendage. The LAA was separated from the LA body by a plane fitted to the ostium on the reference configuration. At each sampled time instant, only Eulerian fluid voxels inside the corresponding ROI were included. For either ROI, the emptying fraction (EF) was calculated from the maximum and minimum volumes during the cardiac cycle as

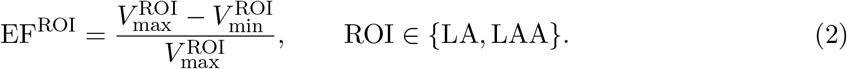

To separate passive from active emptying, the LA EF was further decomposed into a passive (reservoir and conduit) and an active (booster) component as in Telle *et al.* [24]

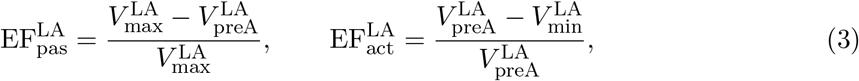

where 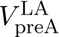 is the LA volume at the onset of electrical activation.

Unless stated otherwise, hemodynamic quantities were sampled at 40 equidistant instants per cardiac cycle during the final 12 CFD cycles (cycles 21–32), yielding 480 sampled instants per case.

For periodic quantities such as chamber volume, flow rate, and kinetic energy, t/T denotes phase within the cardiac cycle (0 ≤ t/T < 1), with t = 0 corresponding to the onset of atrial electrical activation. For residence-time analyses spanning multiple cycles, t/T denotes cumulative normalized simulation time. Specific kinetic energy was calculated at each fluid voxel as 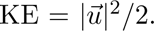 For scalar case-level comparisons, we used the non-dimensional kinetic energy introduced by Gonzalo *et al.* [22]:

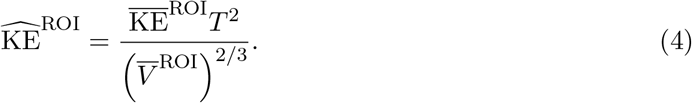

Here, 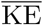 is the time- and space-averaged KE, 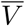 the time-averaged volume, and T the duration of a cardiac cycle. For a scalar residence-time metric, voxel-wise T*_R_* values within each ROI were collected over the 40×12 sampled time points. The upper tail of the resulting pooled distribution was summarized by its 90^th^ percentile to characterize prolonged blood residence while reducing sensitivity to isolated extreme values.

## 3 Results

In the following, we present detailed results from the EM and CFD simulations with particular emphasis on KE and T*_R_*. The first part focuses on cases with fibrosis applied throughout the atrium, referred to as global-fibrosis cases. The second part focuses on cases in which fibrotic tissue was restricted to the LAA, called LAA-fibrosis cases.

### 3.1 Effect of Globally Distributed Fibrosis

Three spatially distinct fibrosis distribution patterns were generated, denoted A, B, and C. Each pattern was applied at two burden levels, 15.6% and 31.2% of atrial tissue volume. This yielded six global-fibrosis cases, which are shown in Figure 2 together with the reference case. Cases are named by a letter denoting the spatial fibrosis pattern and a number denoting the global fibrotic burden. Cases with 15.6% fibrotic tissue (A15, B15, C15) are referred to as low-fibrosis cases, whereas cases with 31.2% fibrotic tissue (A31, B31, C31) are called high-fibrosis cases.

**Fig. 2.**
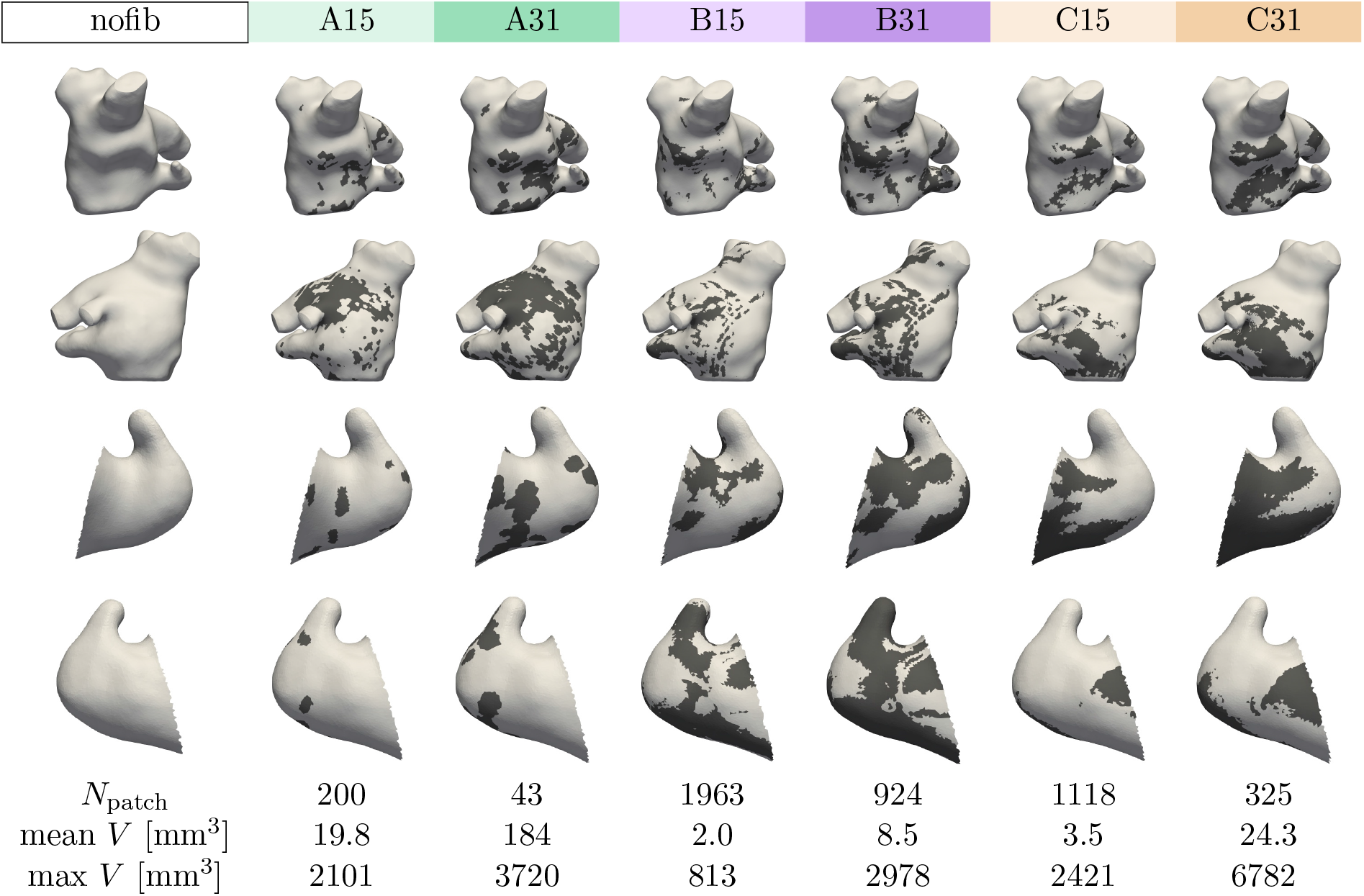
Spatial fibrosis distributions for the global-fibrosis cases. The first and second rows show anterior and posterior views of the whole LA, respectively; the third and fourth rows show two views of the isolated LAA. Dark regions denote fibrotic tissue. The last three rows report the number of fibrotic patches and their mean and maximum volume, respectively, which are determined as described in § 2.3

All three patterns were patchy and spanned the entire LA body, but they differed in both their predominant locations and their fragmentation, as shown and quantified in Figure 2. Pattern A was mainly localized at the LA roof near the left PVs, with limited extension into the LAA, and consisted of comparatively few, large patches. Pattern B was more diffusely patchy throughout the LA body and had the densest fibrotic distribution within the LAA. At baseline burden, it contained approximately ten times as many patches as pattern A, with a maximum patch volume approximately 38% of that in pattern A. Pattern C was concentrated toward the MV in the LA body and, within the LAA, toward the LAA base and center, with fragmentation intermediate between patterns A and B. Doubling the burden reduced the number of patches by factors of approximately two to five, while increasing their mean and maximum volumes. These changes indicate that additional fibrotic tissue predominantly merged with existing patches rather than forming new ones.

#### 3.1.1 Electromechanical Simulation Results

Figure 3 compares representative EM results of nofib with A31, the high-burden case for fibrosis pattern A. Relative to nofib, A31 showed only small, spatially heterogeneous changes in activation time (Fig. 3a).

**Fig. 3.**
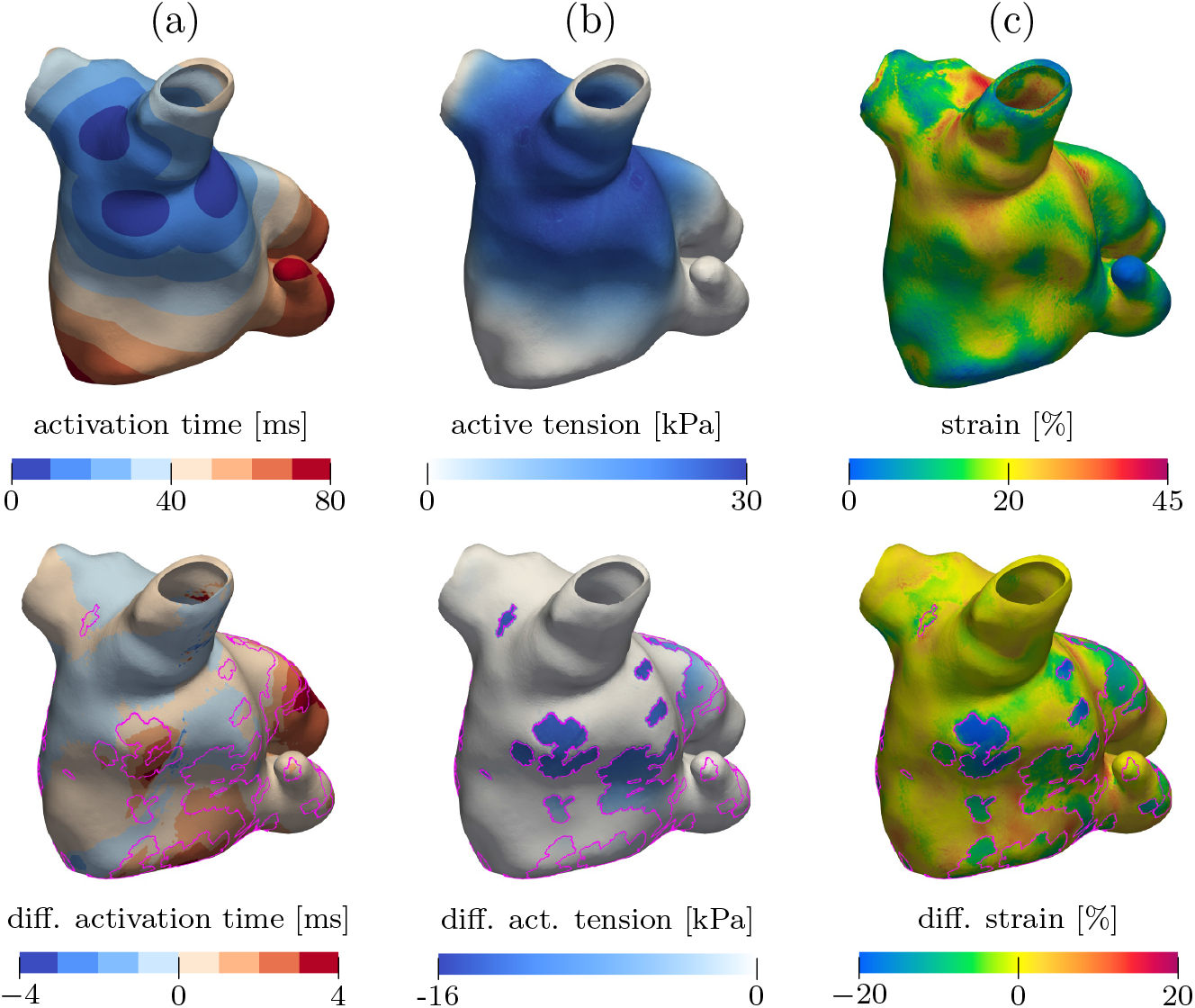
Representative EM results for nofib and A31. (a) Electrical activation time. (b, c) Spatial distributions at 70 ms after excitation of (b) active cellular tension and (c) first principal strain. The first row shows nofib; the second row shows the difference of A31 relative to nofib. Fibrotic regions in A31 are outlined in pink

At 70 ms after excitation, active cellular tension was lower in A31 than in nofib, with the reductions concentrated primarily within the fibrotic regions, consistent with the scaling applied in these regions (Fig. 3b). The first principal strain showed a less localized response. Although the largest reductions coincided with fibrotic regions, differences extended into non-fibrotic myocardium, where both increases and decreases relative to nofib were observed (Fig. 3c).

#### 3.1.2 Fibrosis-Induced Changes in Chamber Function, Volume Dynamics, and Flow Rates

Increasing fibrosis burden enlarged the LA and LAA and reduced their EF (Tab. 2), whereas left ventricular stroke volume (LVSV) remained nearly unchanged across cases. Both components of LA emptying were affected: passive LA EF decreased from 17% to 14%, and active LA EF from 22% to 18–19%.

**Tab. 2.** Anatomical and functional chamber characteristics. Fibrotic burdens refer to the EM mesh; volumes and EF were computed from the CFD ROI voxels (Eqs. 2 and 3). Cell colors identify the cases throughout the manuscript

| case |  | nofib | A15 | A31 | B15 | B31 | C15 | C31 |
| --- | --- | --- | --- | --- | --- | --- | --- | --- |
| fibrotic burden [%] | LA | 0 | 15.6 | 31.2 | 15.6 | 31.2 | 15.6 | 31.2 |
|  | LAA | 0 | 6.9 | 22.0 | 37.4 | 67.5 | 15.3 | 36.5 |
| mean $V$ [mL] | LA | 122 | 132 | 137 | 132 | 138 | 132 | 138 |
|  | LAA | 4.0 | 4.3 | 4.5 | 4.5 | 4.7 | 4.5 | 4.7 |
| EF [%] | LA, total | 36 | 32 | 30 | 33 | 30 | 33 | 31 |
|  | LA, passive | 17 | 15 | 14 | 16 | 14 | 15 | 14 |
|  | LA, active | 22 | 20 | 18 | 20 | 19 | 21 | 19 |
|  | LAA | 44 | 43 | 38 | 34 | 28 | 37 | 31 |
| LVS<sub>V</sub> [mL] |  | 83.2 | 82.9 | 82.5 | 82.9 | 82.7 | 83.2 | 83.1 |

Atrial contractile work decreased with increasing fibrosis burden: both the maximum A-loop pressure and the area enclosed by the A-loop were smaller in A15 than in nofib, and smaller still in A31 (Fig. 4a). Across all patterns, EF decreased with increasing burden in both the LA and LAA (Fig. 4b).

**Fig. 4.**
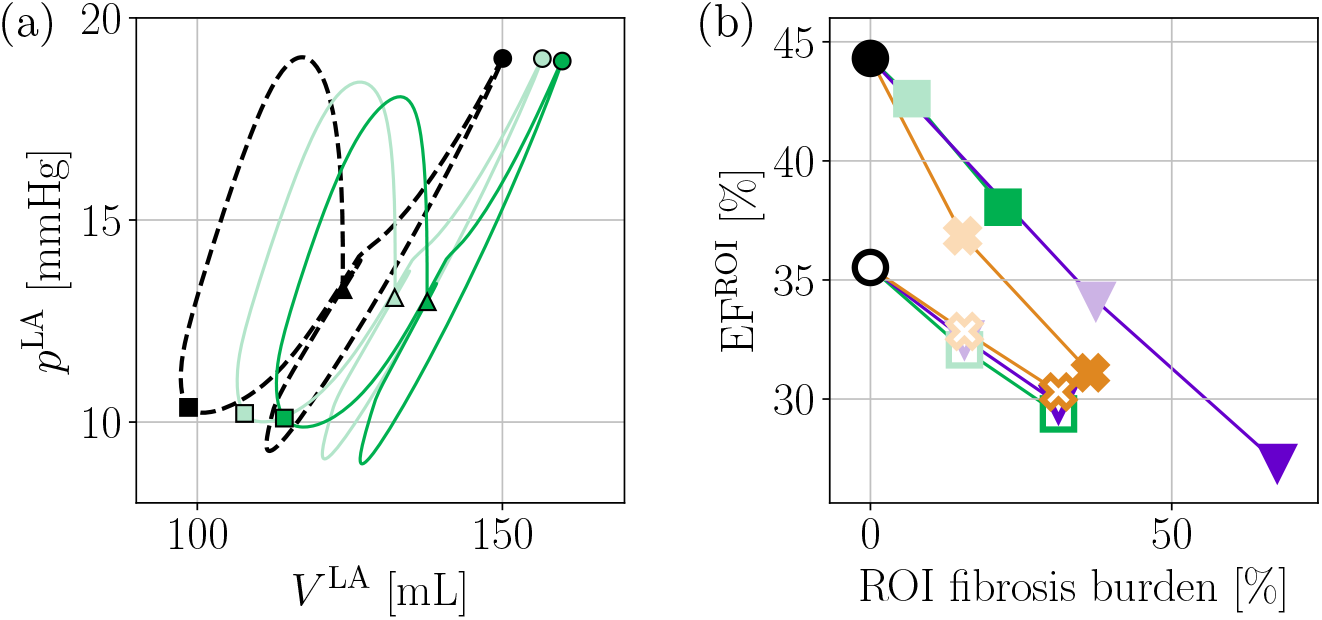
(a) Left atrial pressure–volume relation for 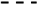 nofib, 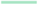 A15, and 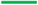 A31. Markers indicate 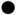 MV opening, 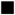 MV closure, and 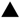 the instant corresponding to Fig. 3b,c. (b) Emptying fraction versus region-specific fibrosis burden in the LA (unfilled markers) and LAA (filled markers). Marker shapes denote patterns 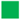 A, 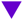 B, and 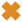 C, with 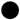 nofib shown for reference

To compare mass exchange among the global-fibrosis cases, we plot the transmitral outflow rate obtained from the 0D circulatory model in Figure 5a. The corresponding CFD flow rate follows the same waveform, as the prescribed atrial wall motion and PV flow rates determine transmitral outflow through mass conservation. It had two distinct peaks per cycle: The A-wave started at t/T ≈ 0.05 and reached its maximum at t/T ≈ 0.125. After atrial expansion, during which Q^MV^ = 0 mL/s, the E-wave started at t/T ≈ 0.56 and reached its maximum at t/T ≈ 0.65. As highlighted in the insets of Figure 5a, both peaks were lower in the fibrotic cases than in nofib. At matched global burden, the A-wave peak depended on the spatial fibrosis pattern, whereas the E-wave peak varied little across patterns.

**Fig. 5.**
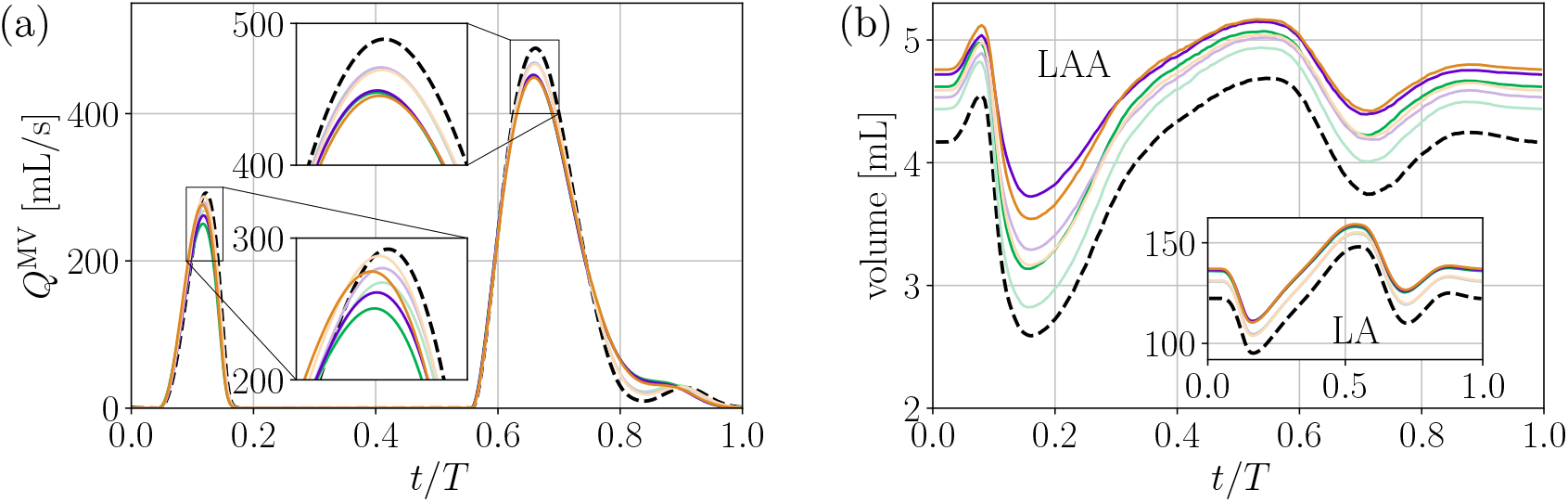
(a) Evolution of transmitral outflow rate and (b) LAA and (inset) LA volume over one cardiac cycle for the 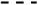 non-fibrotic case and the fibrotic cases 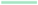 A15, 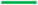 A31, 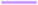 B15, 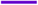 B31, 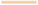 C15, and 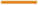 C31. Insets in (a) show zoomed views of the A-wave and E-wave peaks

Figure 5b shows the temporal evolution of LAA volume, with LA volume shown in the inset. The LA volume curves were approximately parallel, with an upward offset that increased with global fibrosis burden. The LAA curves retained a similar overall shape but were not strictly parallel: fibrosis had its most pronounced effect near the minimum at t/T ≈ 0.15 and raised the minimum volume as burden increased.

#### 3.1.3 Impact of Global Fibrosis on Flow and Residence Time Fields

Figure 6 shows flow fields for three representative cases, nofib, A15, and B31, at the peak A-wave and E-wave instants. These cases were selected to cover a broad range of LAA function (Tab. 2 and Fig. 4b). To reduce effects of cycle-to-cycle variability, the figure shows phase-averaged results over the final 12 simulated CFD cycles.

**Fig. 6.**
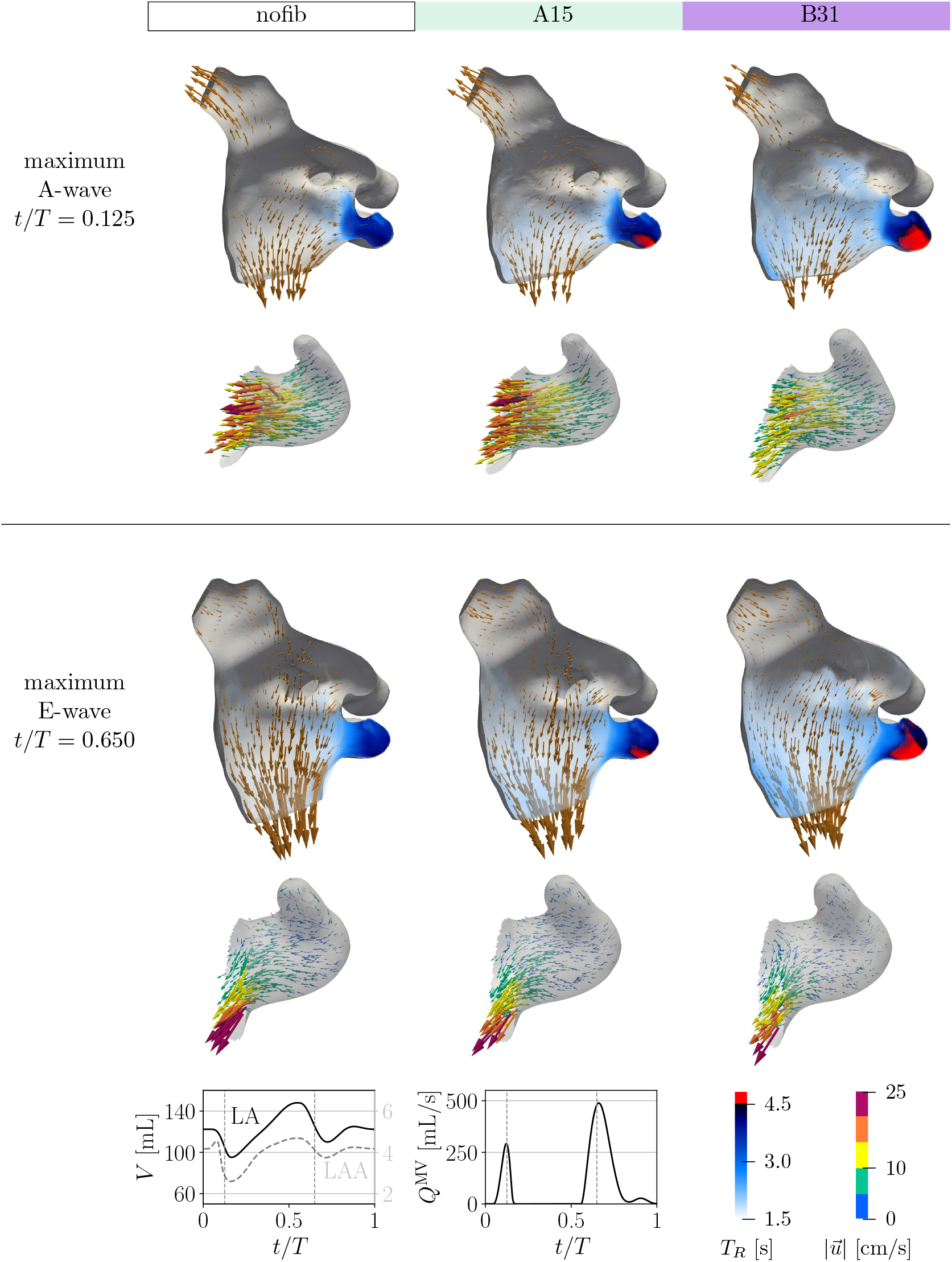
Visualization of phase-averaged flow fields for three representative cases (nofib, A15, B31) at the instants of maximum A-wave and maximum E-wave. Rows 1 and 3: volumetric rendering of residence time in the entire LA with velocity vectors on a planar cross-section. Rows 2 and 4: volumetric velocity vectors within the LAA, colored by velocity magnitude. Bottom panels show the evolution of volume and Q^MV^ for the nofib case, with vertical dashed lines marking the selected time instants

Volumetric renderings of residence time across the entire LA are shown alongside velocity vectors on a planar cross-section that captures the LAA. Only regions with T*_R_* ≥ 1.5 s are displayed for clarity. Additionally, volumetric velocity vectors within the LAA are shown to visualize appendage flow. A localized region of elevated T*_R_* is visible inside and near the LAA. In the nofib case, most of the LA body remained below the lower visualization threshold of T*_R_* = 1.5 s. Compared with nofib, the region exceeding this threshold became larger as fibrosis burden increased, particularly near the LAA orifice and in the near-wall region toward the MV. Similarly, the high-T*_R_* regions exceeding 4.5 s, highlighted in red, grew as the burden increased.

At the A-wave peak, atrial contraction drove flow out of the LA through both the MV and PVs in all three cases. In the LAA, nofib exhibited a coherent outflow jet, with the highest velocities concentrated near the ostium. The peak velocity and spatial extent of this jet were lower in A15 and lower still in B31.

During the E-wave, flow entered the LA from the PVs and exited through the MV. Within the LAA, the flow resembled that of a contracting, lid-driven cavity, consistent with the LA chamber serving primarily reservoir and conduit functions during this phase. At the ostium, the outward velocity component decreased relative to the tangential component as fibrosis burden increased. Overall, the LAA velocity magnitudes were lower during the E-wave than during the A-wave in all three cases.

#### 3.1.4 Impact of Global Fibrosis on Blood Kinetic Energy and Residence Time Global Flow Metrics

Figure 7 presents the temporal evolution of the spatially averaged KE and T*_R_* in the LA. The low-fibrosis cases (A15, B15, C15) and high-fibrosis cases (A31, B31, C31) are shown separately. Considering the low cycle-to-cycle variability of the quantities, as illustrated by the insets of Figure 7a,c for the non-fibrotic case, the main panels show one representative cycle. To facilitate comparison, the high-fibrosis envelope is overlaid as a gray band in the low-fibrosis panels (a,c), whereas the low-fibrosis envelope is overlaid in the high-fibrosis panels (b,d).

**Fig. 7.**
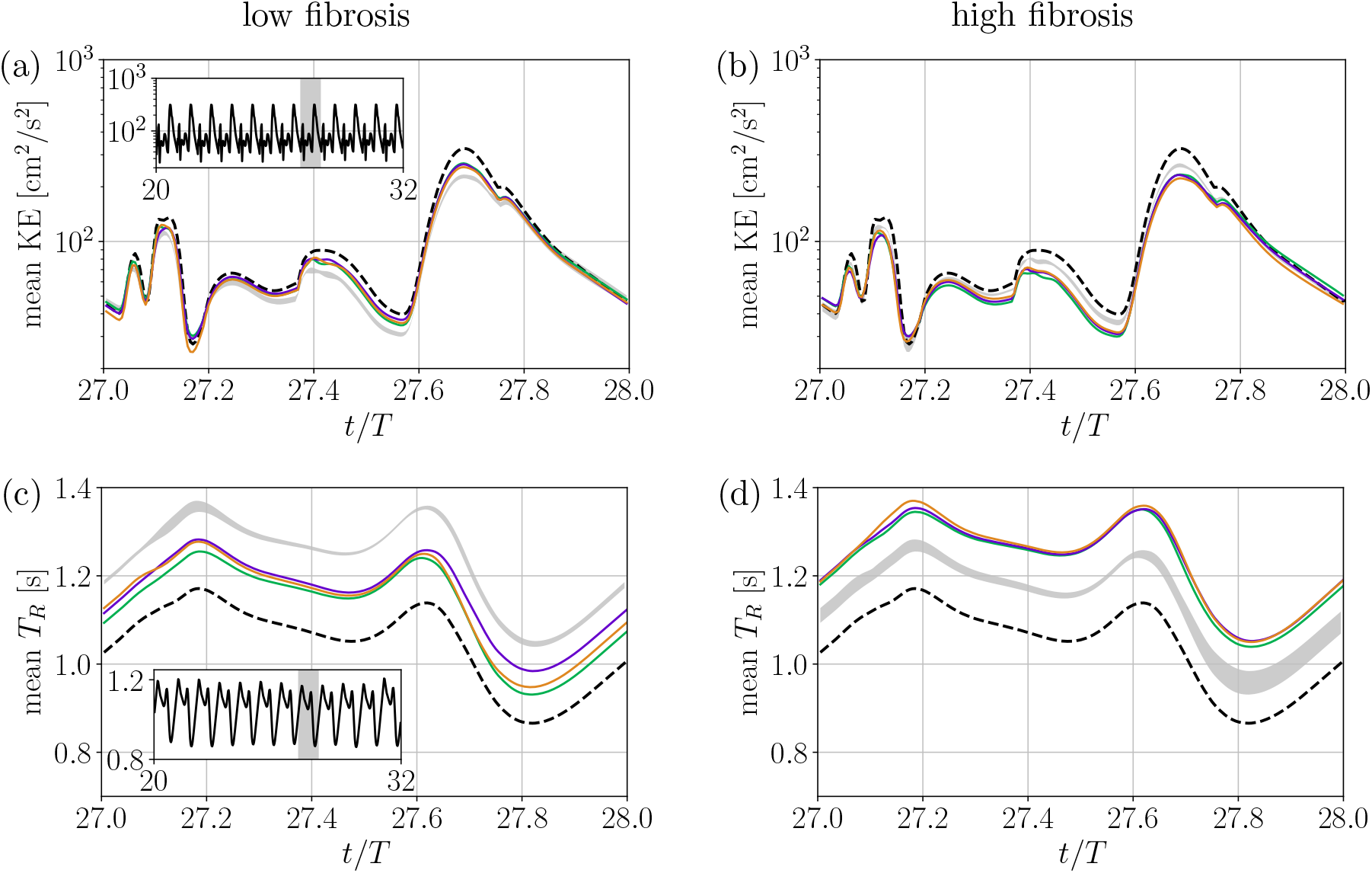
Temporal evolution of the spatially averaged (a, b) specific kinetic energy and (c, d) residence time within the LA of the (a, c) low-fibrosis and (b, d) high-fibrosis cases with patterns 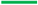 A, 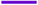 B, and 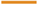 C. The 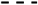 non-fibrotic case is plotted for reference in all panels. The gray bands in the main panels show the envelope of the respective other fibrosis-level group.

The insets in (a, c) show the same quantity over 12 cycles for the non-fibrotic case and the shaded region marks the cycle shown in the main panels The temporal behavior and numerical values of LA KE and T*_R_* align with previous patient-specific simulations driven by prescribing wall motion from time-resolved computed tomography (CT) images [23, 49, 50]. These quantities evolved quasi-periodically from cycle to cycle, with recurrent peaks and valleys associated with atrial contraction, PV inflow, and mitral outflow.

The spatially averaged KE (Fig. 7a,b) fluctuated along the cardiac cycle, with primary and secondary peaks coinciding with the E-wave (t/T ≈ 0.65) and the A-wave (t/T ≈ 0.125). Mean KE in the low-fibrosis cases remained below nofib while largely preserving its waveform, and was further reduced in the high-fibrosis cases throughout the cycle. For both groups, the greatest separation from nofib occurred at the E-wave peak. The low- and high-fibrosis envelopes were narrow and showed little to no overlap.

Mean T*_R_* (Fig. 7c,d) rose during the A-wave, decreased during the subsequent atrial expansion, and then reached a second local maximum in the early E-wave, before decreasing sharply. The nofib case had the lowest mean T*_R_* throughout the cycle. Fibrotic cases showed elevated mean T*_R_*, with higher values in the high-fibrosis cases than in the low-fibrosis cases. As for mean KE, the low- and high-fibrosis envelopes of T*_R_* were narrow and largely non-overlapping.

#### LAA Flow Metrics

The corresponding KE and T*_R_* evolutions within the LAA are shown in Figure 8. In contrast to the LA, where the dominant KE peak occurred during the E-wave, mean KE in the LAA reached its dominant peak during the A-wave. The A-wave peak magnitude was clearly reduced with increasing fibrosis burden. The subsequent E-wave produced only a smaller secondary local maximum.

**Fig. 8.**
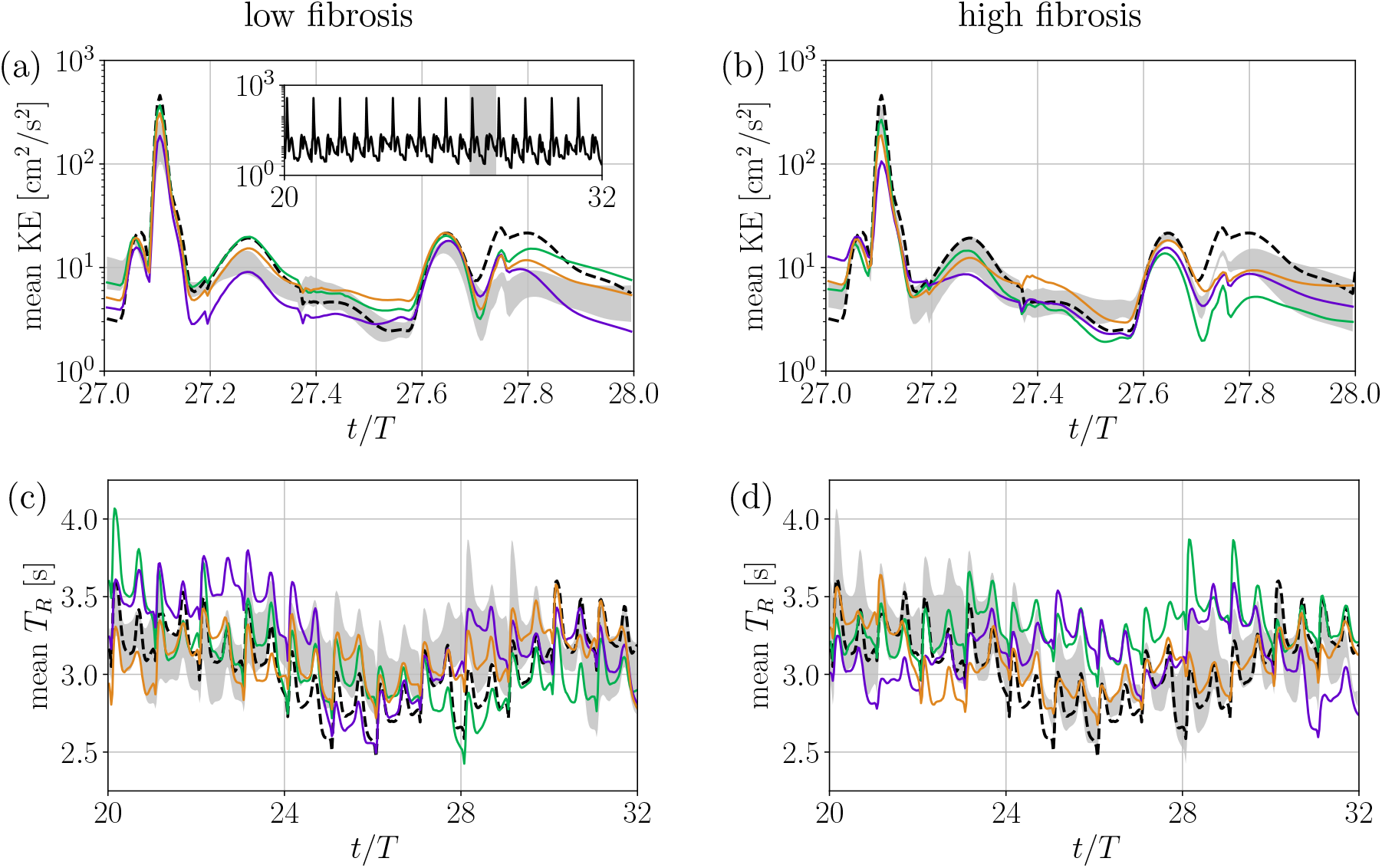
Temporal evolution of the spatially averaged (a,b) specific kinetic energy and (c,d) time-resolved residence time within the LAA, for low-fibrosis (a,c) and high-fibrosis (b,d) cases with patterns 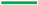 A, 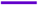 B, and 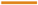 C. The 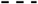 non-fibrotic case is plotted for reference in all panels. The gray bands in the main panels show the envelope of the respective other fibrosis-level group. Kinetic energy is shown for one representative cycle (inset in (a): nofib case over all 12 cycles); residence time is shown over all 12 cycles due to pronounced cycle-to-cycle variability

Between the A-wave and the E-wave, the KE trajectories diverged between cases, with no clear ordering of the cases. For nofib, the magnitudes of both peaks were similar across the 12 cycles shown in the inset of Figure 8a, whereas the trajectories between the peaks exhibited greater cycle-to-cycle variability. Hence, the single-cycle comparison within these phases should not be over-interpreted as a robust burden effect without further verification.

In contrast to the LA, T*_R_* in the LAA exhibited pronounced cycle-to-cycle variability throughout all cardiac phases. We therefore report T*_R_* for the LAA over all 12 cycles in Figure 8c,d. Over this interval, mean T*_R_* fluctuated irregularly, with no case consistently highest or lowest, and the low- and high-fibrosis envelopes overlapping almost entirely throughout the cycles.

#### 3.1.5 Relation between Hemodynamic Metrics and Chamber Emptying Fraction

Figure 9 shows how non-dimensional kinetic energy 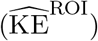 and the 90^th^ percentile of residence time varied with EF. Values are displayed separately for the LA and LAA in each simulation case. 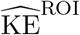 increased with EF^ROI^ (Fig. 9a). For the LA, this relationship was approximately quadratic and consistent with the trend reported by Gonzalo *et al.* [22].

**Fig. 9.**
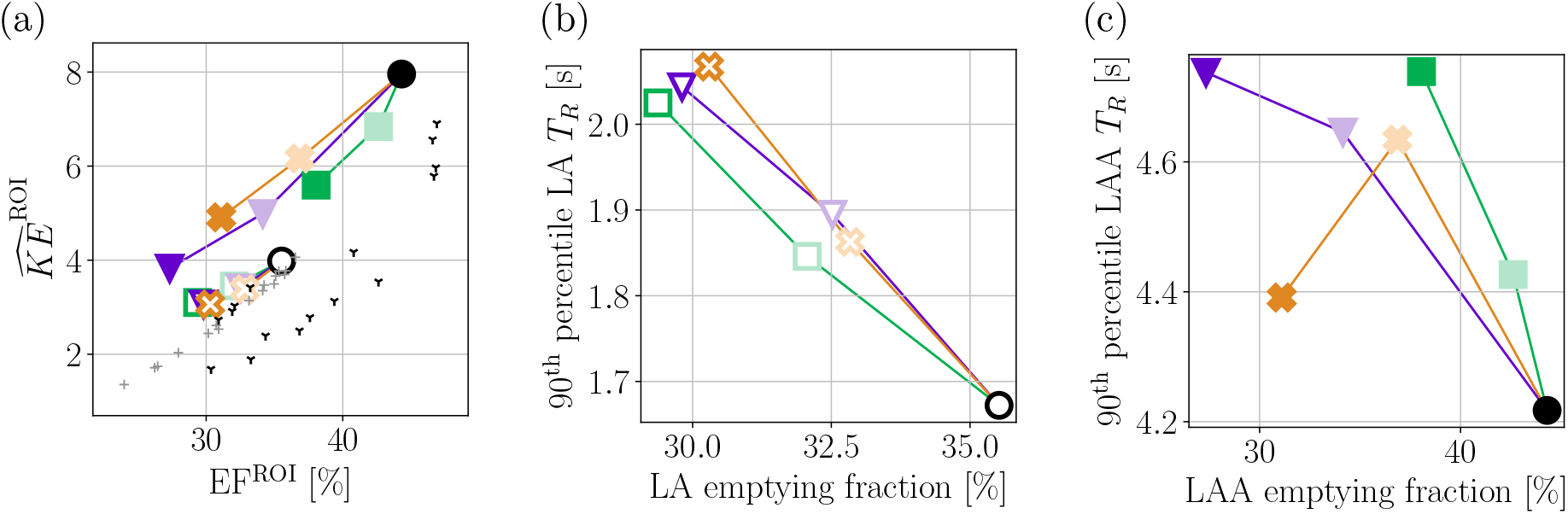
(a) Non-dimensional kinetic energy versus region-specific emptying fraction. (b, c) 90^th^ percentile of residence time versus region-specific emptying fraction in the (b) LA and (c) LAA. Unfilled symbols correspond to quantities of the LA, while filled symbols represent the LAA. Colors and symbols are consistent with Figure 4. In (a), we additionally plot results from previous simulations presented in Gonzalo *et al.* [22] for the 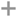 LA and the 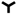 LAA

Compared with the booster-pump simulations of Gonzalo *et al.* [22], the present full-cycle simulations followed a similar trend but yielded higher 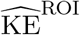 values. This difference may reflect the contribution of E-wave flow, which was not included in the earlier simulations. Within the LAA, 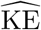 also generally increased with EF, but the points were more dispersed than for the LA and reached higher values than those reported by Gonzalo *et al.* [22].

In the LA, the 90^th^ percentile of T*_R_* was higher at lower EF across all patterns (Fig. 9b). For the LAA, lower EF was likewise accompanied by higher T*_R_* for most cases, but C31 deviated from this trend (Fig. 9c).

### 3.2 Effect of Fibrosis in the Appendage

To isolate the influence of impaired LAA wall motion on local flow, we performed six additional simulations with fibrosis confined to LAA tissue, as shown in Figure 10. The cases are named using the prefix “LAA-”, followed by the fibrosis pattern (A or B) and local fibrotic burden (10%, 25%, or 50%). The nofib model serves as the reference. Across the LAA-fibrosis cases, global LA EF differed from the nofib value by less than 1% in relative terms.

**Fig. 10.**
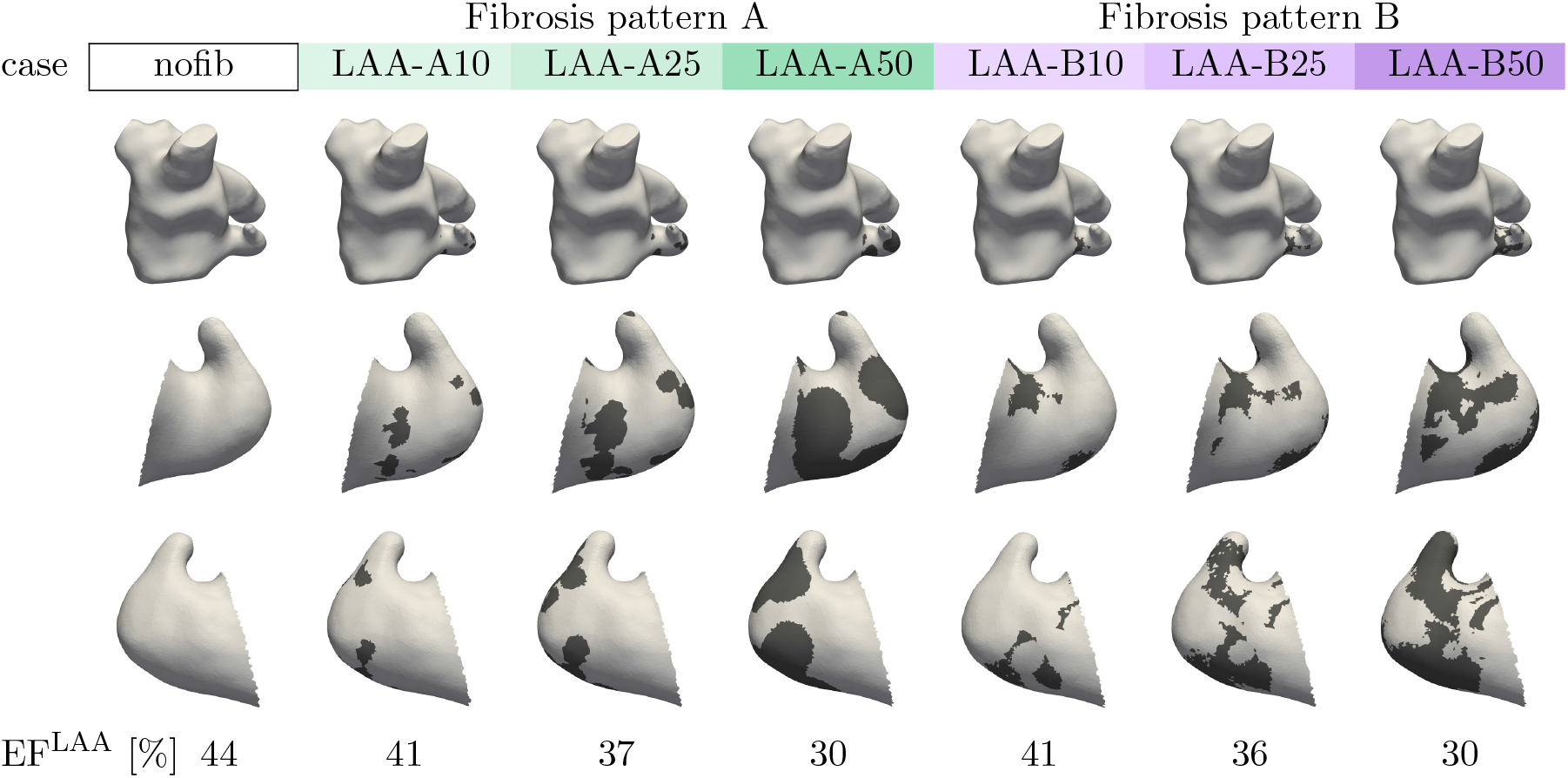
Fibrosis distributions and LAA EF for the LAA-fibrosis cases. The top row shows anterior views of the whole LA; the middle and bottom rows show two views of the isolated LAA. Dark regions denote fibrotic tissue, and the values below the bottom row report LAA EF

Non-dimensional LAA kinetic energy decreased systematically with decreasing LAA EF (Fig. 11a). Patterns A and B followed the same relationship despite their different spatial fibrosis distributions. In contrast, for the 90th percentile of LAA T*_R_* (Fig. 11b), the two fibrosis patterns did not collapse onto a common relationship with LAA EF. Moreover, the variation of the 90^th^ percentile of T*_R_* with LAA EF was not strictly monotonic.

**Fig. 11.**
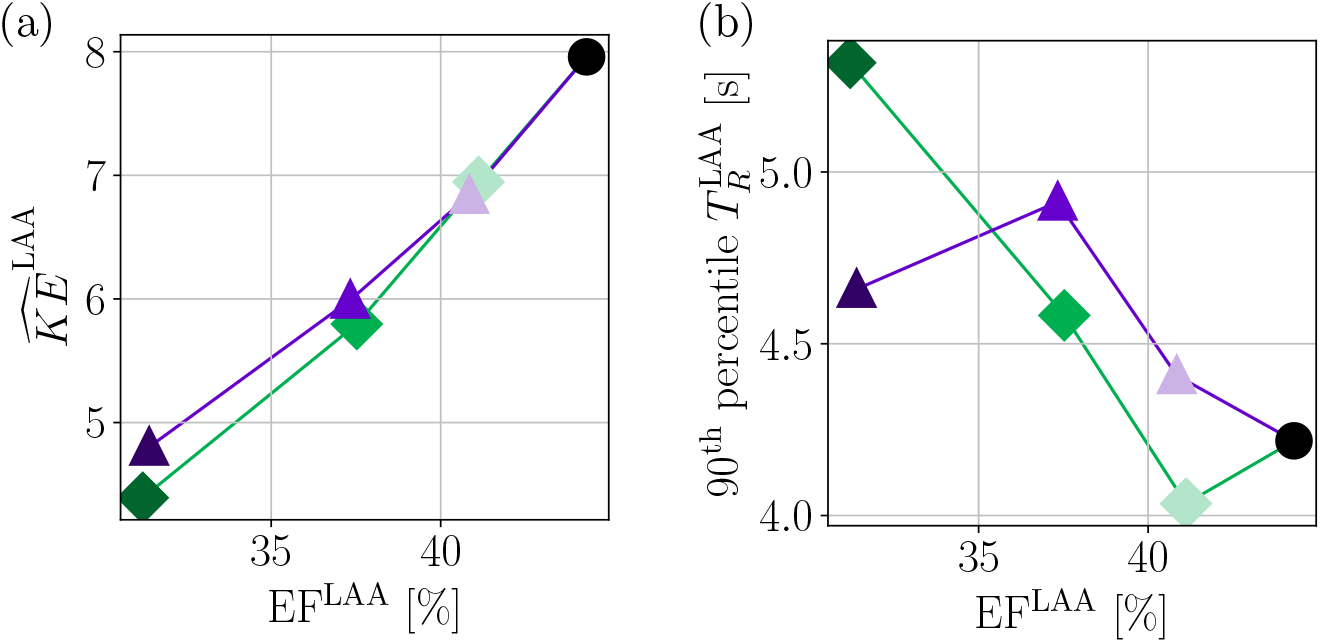
(a) Non-dimensional LAA kinetic energy and (b) 90^th^ percentile of LAA residence time versus LAA emptying fraction for the fibrosis patterns 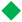 A, 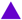 B, and the 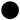 non-fibrotic reference case

## 4 Discussion

Atrial fibrosis is associated with impaired atrial function and thromboembolic risk, motivating interest in LGE–MRI fibrosis measurements for stroke-risk stratification [3]. However, these clinical associations do not identify the mechanisms linking structural remodeling to blood stasis [15]. Regional heterogeneity and differences in fibrotic patch size and location at comparable global burdens further motivate considering spatial distribution separately from total fibrosis burden [51, 52].

Gonzalo *et al.* [22] showed that fibrosis-induced mechanical impairment alters flow throughout the atrium, but their focus on booster-pump function did not allow quantification of physiological long-term transport. The present study extends this approach to full-cycle, multi-beat simulations on a common atrial anatomy. Complementary global- and LAA-fibrosis cases help distinguish global and local functional changes, and allow the effects of burden and spatial distribution to be examined separately. The findings indicate that the spatial distribution of contractile impairment contributes information about LAA transport beyond what is captured by bulk emptying alone.

### 4.1 Whole-Atrium Flow and Transport

Across the global-fibrosis cases, global LA hemodynamic metrics varied more consistently with fibrosis burden than with spatial pattern. Increasing burden impaired atrial emptying, weakened both the E- and A-wave transmitral outflow jets, reduced kinetic energy, and increased residence time. The approximately quadratic trend between non-dimensional 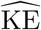 and LA EF is consistent with the empirical relationship reported by Gonzalo *et al.* [22]. A simple mass-conservation scaling argument supports the quadratic trend. We take the change in LA volume, LA cross-sectional area, and flow rate to scale as ΔV = EF V, A ∼ V ^2*/*3^, and Q ∼ ΔV/T, respectively, where V is a characteristic LA volume and T is the cardiac period. Consequently, 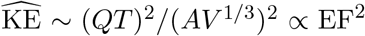 Note that the slightly higher values obtained here compared to Gonzalo *et al.* [22] may reflect our consideration of the full cardiac cycle, including the E-wave. We also found LA residence time to depend almost exclusively on LA EF, regardless of fibrosis distribution. Similarly, Gonzalo *et al.* [49] related LA residence time to EF as T*_R_* ∼ T (EF + V_conduit_/V)*^−^*^1^.

The insensitivity to spatial pattern held despite substantial differences in fibrosis topology: at baseline burden, pattern B contained approximately ten times as many fibrotic patches as pattern A (Fig. 2), yet LA EF, KE, and T*_R_* were nearly identical. At the whole-atrium scale, the hemodynamic response therefore appears to depend primarily on the amount of impaired tissue, with little sensitivity to its spatial concentration or fragmentation across the patterns examined.

A notable outcome of the full-cycle setup is that the fibrosis-induced reduction in LA KE was largest during the E-wave rather than during atrial contraction. Because fibrotic tissue was modeled with increased passive stiffness in addition to reduced active tension, fibrosis is expected to impair not only the booster function but also the LA reservoir and conduit function, consistent with the reduced passive EF (Tab. 2). This finding agrees with clinical strain-imaging studies that associated more extensive LGE with impaired reservoir and conduit strain, not only with impaired booster strain [6–8]. Together, these findings suggest that EF provides a useful compact descriptor of the hemodynamic effects of fibrosis at the global LA scale.

Separate sensitivity studies provide context for this robust chamber-scale response. These studies found that residence time in the LA body changed little when blood rheology [49] or PV inflow split ratios were varied [50]. Both studies related this behavior to the dominant PV–MV conduit flow being constrained by global chamber mass conservation. Similarly, global LA metrics were less sensitive to temporal wall-motion sampling than LAA metrics [42].

### 4.2 Appendage Stasis Is Multifactorial and Fibrosis-Dependent

The effects of fibrosis on appendage function and stasis were less systematic than those observed at the whole-LA scale. In the more realistic global-fibrosis cases, which used whole-atrial fibrosis patterns derived from LGE data, increasing fibrosis burden reduced LAA EF. However, this reduction did not translate monotonically into increased residence time, suggesting that bulk appendage emptying alone does not explain prolonged blood retention. Moreover, the contribution of local appendage impairment could not be isolated because LA and LAA EF changed simultaneously.

The virtual LAA-fibrosis cases reduced changes in global LA function while varying local appendage impairment. Across these cases, the non-dimensional LAA KE showed a systematic relationship with LAA EF, with different spatial patterns following essentially the same trend. This supports LAA EF as a useful functional descriptor of non-dimensional LAA KE, as in the LA body. However, residence time did not show the same collapse and exhibited non-monotonic trends that varied across fibrosis patterns.

The different residence-time trends at similar LAA EF levels highlight the distinction between bulk emptying and replacement of retained blood. Previous clinical and computational studies show that LAA anatomical characteristics [53–57], variations in the PV inflow jets [50, 58–60], and blood rheology [49, 61] also change atrial flow patterns and LA–LAA exchange, and thereby influence appendage washout.

These findings connect to the clinical motivation for patient-specific assessment of atrial thromboembolic risk. Conventional scores such as CHA_2_DS_2_-VASc largely reflect general vascular risk factors and can predict thromboembolic events even in populations without diagnosed AF [2, 62], emphasizing that their information is not specific to an atrial transport mechanism. The present study forms part of an ongoing effort to translate patient-specific atrial hemodynamics into mechanistic thrombotic-risk metrics. This effort builds on associations between LAA residence time and prior thromboembolic history [23], and on patient-specific coagulation modeling linking hemodynamics to thrombin accumulation [63].

Because LAA EF did not fully explain the upper tail of LAA residence time in these simulations, fibrosis pattern may contain mechanistically relevant information beyond fibrosis burden or bulk functional indices. Whether this information improves patient-level thromboembolic risk stratification will require validation in anatomically diverse cohorts with in vivo flow and clinical outcome data.

### 4.3 Limitations and Future Work

We intentionally used a common atrial geometry for all simulations, which enabled us to isolate the effects of fibrosis burden and spatial distribution from those of anatomy. Fibrosis burden has been found to correlate with atrial anatomical features, especially LA volume, but this correlation seems to be modest and inconsistent across studies [64–66]. Future work should examine the observed trends across a broader range of anatomies and fibrosis distributions. Given the small number of cases per pattern in the present study, these trends were assessed descriptively rather than through formal statistical testing, and should be interpreted accordingly.

Fibrotic remodeling was represented using discrete non-fibrotic and fibrotic tissue classes, with fixed changes in passive stiffness, active tension, and conduction velocity. This binary treatment is common in image-based atrial models [24, 67] and improves on earlier models that represented fibrosis as a uniform stiffness increase [68]. However, this work did not include any changes to ion channel conductances [15, 24, 67]. Inclusion of these, especially reduced L-type calcium conductance, would further reduce the booster function [24]. Furthermore, the adopted scaling factors remain uncertain because quantitative relationships between local LGE intensity or fibrosis fraction and tissue-scale electromechanical properties are lacking. Consequently, our model did not explicitly capture graded or microscale heterogeneity within LGE-defined fibrotic regions or diffuse interstitial remodeling [69] of tissue classified as non-fibrotic, nor did it directly represent the consequences of endomysial fibrosis, which is independently associated with AF [70]. As techniques used to image the diseased LA continue to improve, future work may be poised to incorporate an even more robust representation of the substrate underlying atrial cardiomyopathy.

The imaging-derived spatial patterns are themselves uncertain. Different acquisition and analysis protocols can yield substantially different fibrosis estimates [71–73]. The spatial resolution of LGE–MRI can also be comparable to atrial wall thickness, limiting the resolution of intramural fibrosis [52]. These limitations qualify how precisely an imaging-derived pattern can represent the underlying distribution of contractile impairment. Future work in this space should strive toward representations of the LA wall incorporating more detailed (and validated) image-based reconstruction of the transmural fibrosis pattern, as well as extension to models incorporating heterogeneous wall thickness.

Blood was modeled as a Newtonian fluid with constant viscosity, neglecting the thixotropic and shear-thinning behavior. This simplification may affect flow and residence time in low-shear regions, particularly within the LAA [50, 60]. Future studies should therefore incorporate non-Newtonian blood rheology.

We used a one-way coupling approach to transfer wall motion from the EM model to the CFD model. Although the EM model received pressure loading from the coupled circulation model, spatially resolved CFD loads were not fed back to the tissue. The consequences of this modeling choice should be assessed in future work.

Fibrosis has been proposed as a potential promoter of atrial thrombosis through impaired contraction and blood stasis, as well as through tissue injury affecting coagulation [4]. The present study addressed only the first pathway, using residence time to characterize the persistence of blood within the LA and LAA. While T*_R_* does not directly model coagulation or thrombus formation, it captures the time available for coagulation reactions to evolve and therefore constitutes an important hemodynamic input to thrombogenesis [63, 74, 75]. However, its ability to represent localized triggers of the extrinsic cascade associated with disturbed wall shear stress and fibrotic tissue injury needs to be examined in future studies.

Beyond these modeling aspects, the simulation-derived metrics require validation against in vivo measurements, such as transesophageal echocardiographic markers of LAA function and stasis [4, 17] and 4D-flow MRI [76]. Subsequently, larger cohorts are needed to assess whether the regional LGE pattern and fibrosis-informed transport metrics improve thromboembolic risk discrimination beyond atrial structure and function, AF characteristics, and clinical risk scores [2, 51, 53, 62].

## 5 Conclusion

This study combined patient-specific EM simulations with multi-beat CFD simulations to examine how fibrosis burden and its spatial distribution relate to blood transport and residence time in the LA and LAA. Across the global-fibrosis and LAA-fibrosis cases, global LA transport and non-dimensional LAA KE were predominantly associated with fibrosis-induced functional impairment. Within the LAA-fibrosis cases, LAA EF emerged as the relevant functional descriptor of non-dimensional LAA KE, although flow from the LA body and other non-local interactions may still modulate this relationship.

In contrast, LAA residence time was not fully explained by LAA EF alone: the 90^th^ percentile of T*_R_*followed distinct, non-monotonic trends for the two LAA-confined fibrosis patterns, indicating that the spatial distribution of fibrosis also affects LAA transport beyond what is captured by bulk chamber emptying in this model.

## Acronyms

AF: atrial fibrillation
CFD: computational fluid dynamics
CT: computed tomography
EF: emptying fraction
EM: electromechanics
EP: electrophysiology
ESUS: embolic stroke of undetermined source
IBM: immersed boundary method
LA: left atrium
LAA: left atrial appendage
LGE: late gadolinium enhancement
LVSV: left ventricular stroke volume
MRI: magnetic resonance imaging
MV: mitral valve
PV: pulmonary vein
ROI: region of interest
TAWSS: time-averaged wall shear stress
WSS: wall shear stress

## Acknowledgments

This work was funded in part by the US National Institutes of Health (NIH), grant numbers R01HL158667 (NA, PMB, CMA, JCA) and R01HL160024 (JCA), and by the Austrian Science Fund (FWF), grant 10.55776/P37063 (CMA).

Electromechanical simulations were performed using computational resources provided by the Austrian Scientific Computing (ASC) infrastructure.

